# Metabolic state and energy reserves jointly regulate adaptive behavioural control

**DOI:** 10.64898/2026.03.22.713499

**Authors:** Marc Tittgemeyer, Bojana Kuzmanovic, Corina Melzer, Frank Jessen, Klaas E Stephan, Lionel Rigoux

## Abstract

Animals continuously adapt their behaviour to changing physiological needs through flexible regulation of behavioural control. However, how the energetic state shapes the distinct components of this adaptive control in humans remains poorly understood. Here, we tested how fasting-induced energy deficits and stored energy reserves jointly regulate inhibitory control and incentive motivation in healthy participants. Fasting selectively increased impulsivity for food rewards, an effect attenuated by body fat percentage. In contrast, fasting increased effort expenditure across both food and monetary rewards, revealing a domain-general increase in motivation best explained by relative energy deficit—the ratio of energy expended during fasting to stored energy reserves. Neither effect was explained by changes in subjective reward valuation. Although self-report questionnaires captured stable behavioural tendencies, they failed to account for state-dependent behavioural variation. Together, these findings show that inhibitory control and motivation are governed by distinct metabolic principles operating across different timescales, revealing that adaptive behavioural control depends on the energetic significance of metabolic state rather than on fasting alone.

## Introduction

To survive, all animals must continuously adapt their behaviour to environmental constraints and bodily needs ^1-3^. Deciding when and how to act, or how much effort to invest, has different fitness consequences depending on the organism’s energetic state. This adaptive flexibility relies on the dynamic regulation of behavioural control, including inhibitory control and motivation, which determines whether actions are suppressed or vigorously pursued. Yet these core components of behaviour are often conceptualised as stable traits, overlooking how metabolic needs and related internal physiological signals continuously shape behavioural control.

Recent integrative frameworks propose that internal physiological signals continuously reconfigure valuation and action-selection processes to promote adaptive behaviour rather than merely maintaining homeostasis ^4-6^. These proposals resonate with recent teleological perspectives that view the brain’s primary function as the prospective regulation of the organism’s state ^7,8^. Together, these perspectives suggest that behavioural control should be understood as a dynamic control policy that continuously adapts to the organism’s physiological state rather than as the expression of stable behavioural traits ^9^.

Hunger provides a particularly tractable model for investigating adaptive behavioural control ^10^. From an ecological perspective, energy deprivation should bias behavioural control toward strategies that favour food acquisition. The benefit of hunger in motivating eating is evident, as it drives behaviour to satisfy underlying caloric needs. Yet, hunger may influence behaviour beyond food consumption, extending to other behavioural domains, consistent with evidence that internal physiological states induce a widespread reconfiguration of valuation and action-selection circuits rather than selectively modulating food-specific pathways ^4,11-13^. Evidence from model organisms further underscores the dynamic nature of the interplay between physiological state and behaviour: in flies, satiety state modulates the transition from local exploitation to global exploration, revealing a state-dependent reconfiguration of navigational strategies and action patterns ^14,15^.

Consistent with this view, behavioural studies in humans indicate that metabolic state regulates multiple components of behavioural control ^16,17^. Hunger increases acquisition-oriented behaviour even for non-food outcomes without a corresponding increase in subjective valuation, suggesting a domain-general enhancement of motivational drive rather than solely modulating food-specific valuation effects ^18^. Furthermore, short-term fasting has been reported to alter inhibitory control ^19,20^, risk assessment ^21,22^, temporal discounting ^23^, and action vigour ^24,25^, while recent work suggests that energetic state directly influences the willingness to exert effort and allocate energetic resources towards goal-directed actions ^26^. However, the nature and domain (i.e. food vs. non-food oriented) specificity of hunger’s effects are still not well delineated. Reviews of motivational neurobiology emphasise that hunger-related signals interact with multiple competing motivational systems and behavioural priorities, highlighting the need for more integrative approaches that assess behavioural adaptation across domains rather than in isolation ^27^.

One possibility is that fasting induces a domain-general reduction in self-control and an increase in motivation across reward types. Alternatively, fasting may selectively prioritise biologically relevant rewards, such as food, while leaving decision-making for abstract rewards relatively unaffected. Empirical evidence has been limited by a focus on single reward domains, most commonly food, which makes it difficult to distinguish among these accounts. Clarifying whether hunger produces domain-general or domain-specific changes in behaviour is essential to understanding the adaptive logic of state-dependent behavioural control.

At the neural level, the animal studies indicate that metabolic and peptidergic signals associated with hunger, such as insulin, ghrelin, and glucagon-like peptide-1 (GLP-1) ^28^, modulate distributed brain circuits and, in particular, dopaminergic pathways implicated in inhibitory control, motivation, effort expenditure and action selection ^27,29-31^, highlighting the mechanistic relevance of midbrain dopamine pathways in linking metabolic signalling to the control of these behavioural dimensions. Recent studies further support this notion by showing that dopaminergic circuits are also affected by metabolic state in humans ^24,32^. Importantly, the same dopaminergic neurocircuits are involved in both food and non-food reward processing, suggesting that the behavioural consequences of their state-dependent modulation could generalise beyond food reward to other incentive domains ^33,34^. More broadly, converging evidence suggests that metabolic signals can gate the translation of rewards into action, effort, and control policies across species ^4,14,35,36^.

Despite these advances, decisive evidence for state-dependent modulation of the neural circuits underlying food- and non-food-related behaviour in humans remains lacking, underscoring the need to clarify the role of metabolic signalling in adaptive behavioural control.

In addition to transient energetic states, individuals differ markedly in their long-term energy reserves, notably stored in body fat ^37-39^. Consequently, identical fasting periods need not represent equivalent physiological challenges: their behavioural consequences should depend on each individual’s context, determined by available energy reserves. Information about these longer-term energy reserves is conveyed to the brain through endocrine signals, most prominently leptin, a peptide secreted by adipose tissue that directly modulates dopaminergic function and motivational circuits ^40-42^. However, these physiological mechanisms are typically interpreted within trait-based frameworks, in which impulsivity and motivation are treated as stable characteristics that contribute to body composition, rather than as adaptive processes dynamically regulated by energetic state.

From an ecological and allostatic perspective, behavioural responses to fasting should thus depend not only on immediate hunger signals but also on energy reserves and basal energy demands. Lean body mass, as the primary determinant of resting energy expenditure ^43^, constrains available energetic resources such that a short-term energetic challenge may have different adaptive significance across individuals. Accordingly, internal state signals, rather than exerting uniform effects across individuals, should be evaluated in relation to longer-term energetic context when shaping the urgency of action and the allocation of effort ^35,36^. The role of body composition in hunger-dependent behavioural adaptation, however, remains largely unexplored.

To address this gap, we therefore asked whether metabolic state differentially regulates inhibitory control and incentive motivation, and whether these effects depend on longer-term energetic reserves. Because these behavioural functions serve distinct adaptive purposes, they may also be governed by different metabolic principles. To answer this question, we assessed, in a series of experiments with human volunteers, how an acute energy deficit induced by fasting interacts with longer-term energetic constraints, as reflected by body composition, to shape adaptive behavioural control. Specifically, we contrasted two alternative hypotheses. Under a domain-general account, fasting should increase motivation and reduce inhibitory control across reward types, reflecting a global shift in behavioural control. Under a domain-specific account, fasting should selectively prioritise biologically relevant rewards, enhancing motivation and reducing inhibitory control primarily in the context of food while leaving behaviour guided by non-food incentives relatively unaffected. Critically, we further hypothesised that these state-dependent effects are modulated by individual differences in long-term energy reserves and basal energetic demands, such that the behavioural consequences of fasting are influenced by body composition. By jointly manipulating metabolic state and the incentive domain while accounting for individual energetic constraints, this approach aimed at clarifying how short- and long-term energetic states interact to flexibly regulate adaptive behavioural control.

## Results

Based on the hypothesis that metabolic needs adaptively regulate behavioural control, we examined how short- and long-term fluctuations in energy state influence inhibitory control and incentive motivation, contrasting food and monetary outcomes to determine whether these effects are domain-general or food-specific. We invited healthy participants (*n* = 116; **Table 1**) to the lab after an overnight fast. During the testing session, participants had the opportunity to earn food and monetary outcomes in a set of behavioural experiments (see **Fig. 1** and Methods) designed to assess motivation (effort expenditure) and impulsivity (inhibitory control). After completing an auction task that captured the subjective valuation of food vs monetary rewards, participants were allowed to consume their wins (eat the food and pocket the money) before completing various questionnaires related to their drive, impulse control, and food behaviour. In addition to these behavioural and self-report markers, we recorded each participant’s body composition and fasting duration to assess metabolic status.

**Figure 1:**
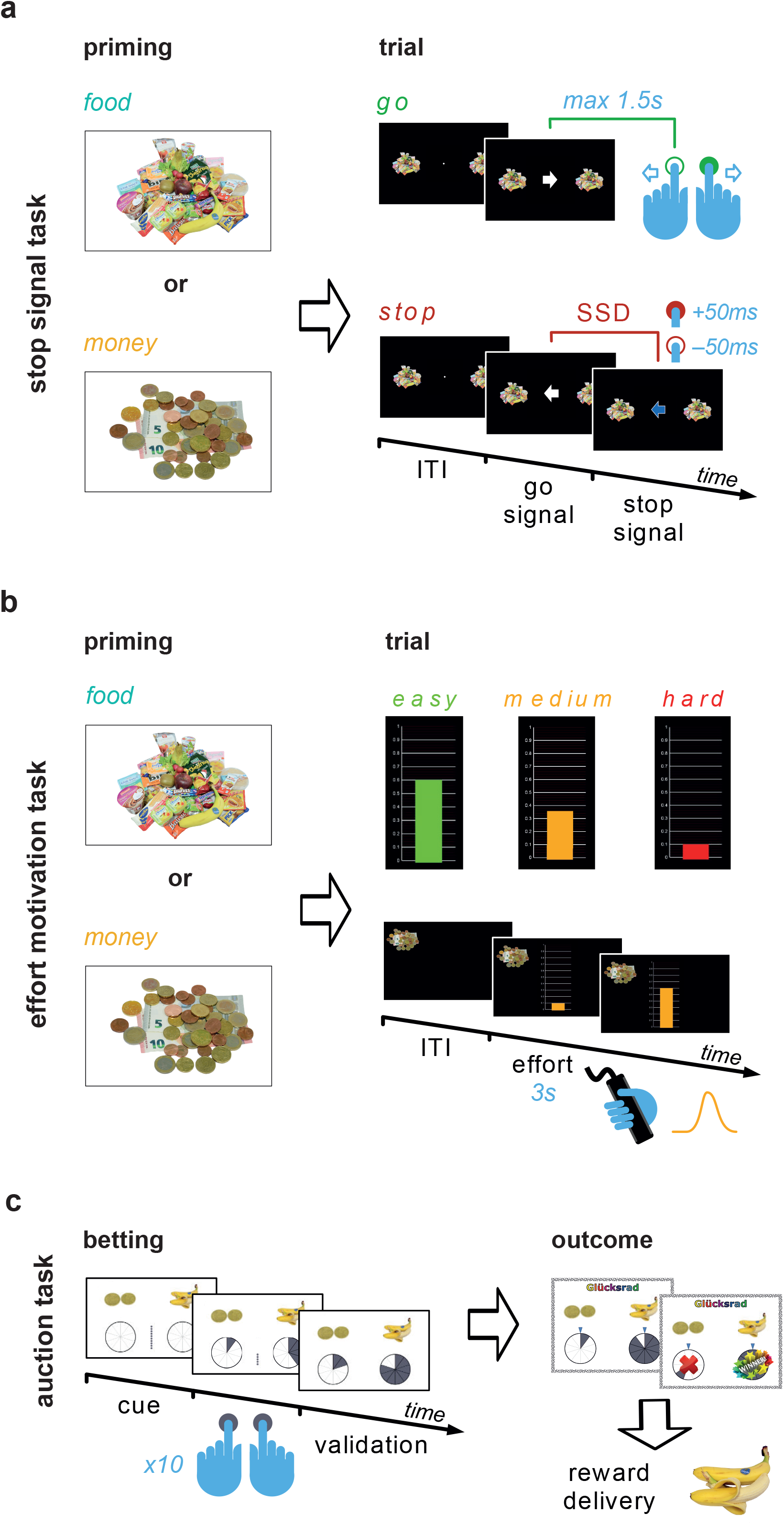
Experimental design. *a) Stop Signal task*. Each block of 96 trials began with a full-screen image depicting the type of reward at stake (food or money). On each trial, a fixation cross first appeared in the centre of the screen, flanked by reward cues. Subsequently, a white arrow was displayed, prompting participants to indicate its orientation via button presses (“go” condition). In 1/4 of the trials, the arrow turned blue after a variable stop signal delay (SSD). In this “stop” condition, participants were instructed to refrain from responding. The SSD was continuously adjusted to induce a 50% chance of correct response inhibition. *b) Effort Motivation task*. Again, each block started with a full-screen display of the reward at stake. On each trial, participants could press a handheld dynamometer to raise the gauge level on the screen, thereby increasing their chances of earning the reward. The gauge’s colour indicated the force required to fill it completely (i.e., the difficulty level). *c: Auction task*. On each trial, participants had to split their bet by placing 10 tokens on either a monetary reward or a food item of equal value (more tokens increased the chance of winning; a maximum of 10 tokens per option). The outcome was displayed once all bets were placed, and both monetary and food (snack) rewards were provided to the participant for consumption.

### Fasting selectively increases impulsivity for food

To quantify variations in impulse control across participants, we adapted the classic stop-signal task ^44^ as the first behavioural task (**Fig. 1a**). Briefly, participants were instructed to press a button or refrain from pressing it in response to go and stop signals, respectively. Trials were organised in alternating blocks, with performance outcomes rewarded with either food or money, as indicated by pictures of the outcomes at the beginning of each block and flanking the go/stop cues. For each participant and each block type, we computed the stop-signal reaction time (SSRT), which captures the time required to stop an ongoing response process; that is, a longer SSRT indicates weaker executive control and therefore more impulsive responding.

A linear mixed-effect model of the SSRT data revealed a strong interaction between fasting duration and reward type (*p* < .001; **Fig. 2a**), which was further modulated by body composition (3-way interaction, *p* = .044). The latter effect was driven by a fasting duration × fat% interaction on SSRT in the food condition (*p* = .040), which could not be observed in the money condition (*p* = .362). To unpack this complex interaction, we calculated the difference in our impulsivity measures between the two conditions *ΔSSRT — SSRT*_food_ *— SSRT*_money_ and used this measure for further analyses.

**Figure 2:**
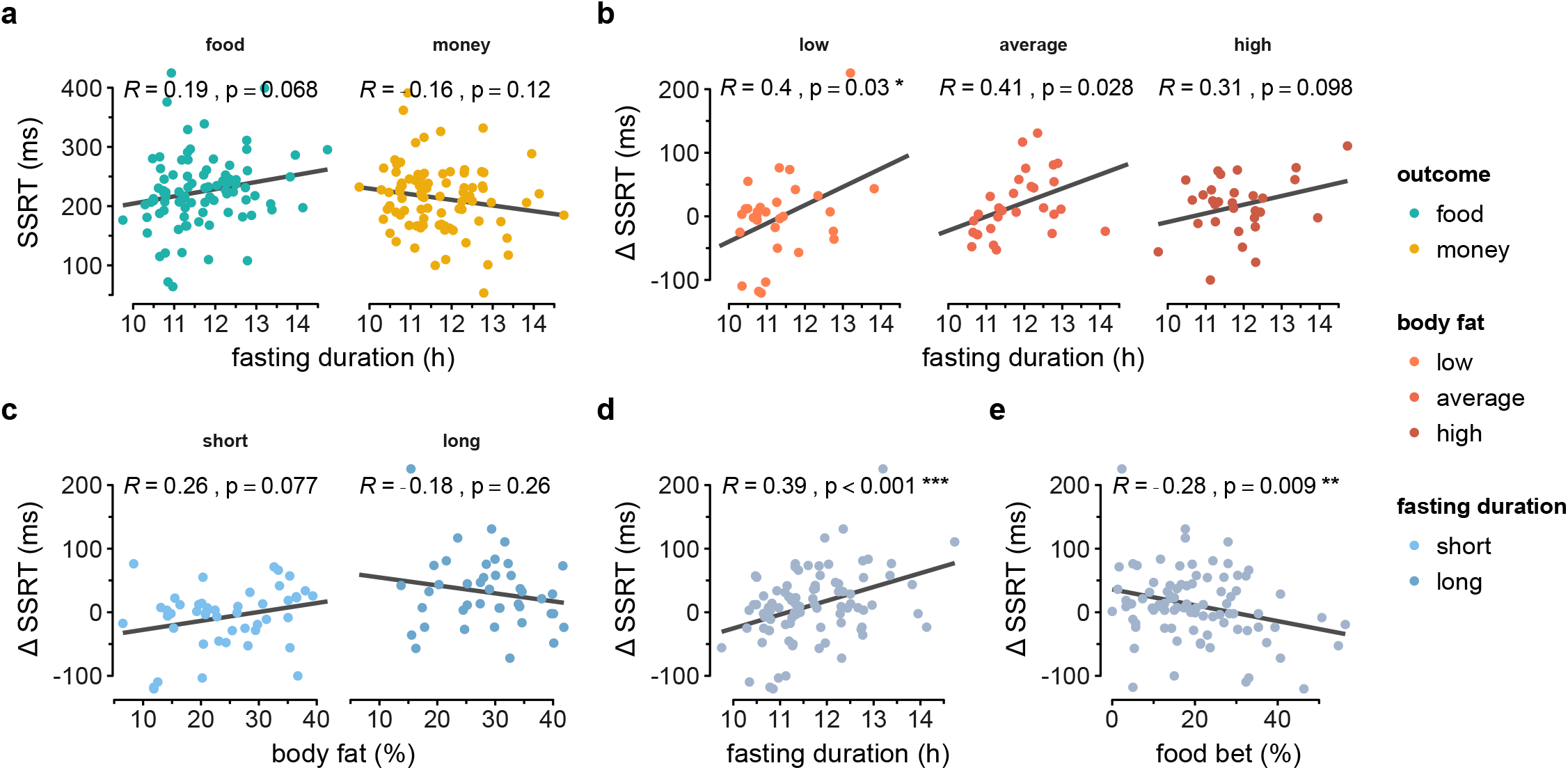
Impulsive behaviour as quantified by the SSRT measured in the stop signal task. *a) SSRT* as a function of fasting duration for food (green) and monetary (yellow) blocs. *b-e)* Relative impulsivity for food relative to money measured by the difference in SSRT between the two conditions (*Δ*SSRT). *b) Δ* SSRT increases with fasting duration, especially for the lowest (*≤* 21%, left, light orange) and central (middle, dark orange), compared to the highest (> 31%, right, red) tercile of body fat percentage. *c) Δ* SSRT increases with body fat percentage for short (< 11.5h, left, light blue) but not for long (> 11.5h, right, dark blue) fasting duration. *d) Δ*SSRT increases with fasting duration (same as in b, collapsed across body composition). *e) Δ*SSRT decreases with larger bets toward food items (as opposed to monetary items) in the auction task. All statistics are Pearson correlations, and the best-fit lines are linear regression fits computed for the data shown in each plot.

A first analysis suggested a fasting × fat% interaction (*p* = .022) which could be understood by the fact that the relative impulsivity for food tended to increase with body fat percentage but fasting mitigated this tendency (short fast: 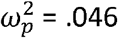; long fast: 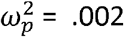; **Fig. 2c**). Strikingly, the impact of fasting on the *ΔSSRT* was stronger for participant with a lower body fat percentage (low body fat: 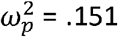; high body fat: 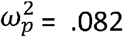; **Fig. 2b**), suggesting that energy reserves modulate the influence of fasting-induced energy deficits. The modulatory effect of body composition, however, reduced to a simpler but highly significant effect of fasting (*p* < .001; **Fig. 2d**) when age and sex were included as confounding factors in the linear model, confirming the strong influence of energy deficit on food-specific impulse control. This follow-up analysis also revealed that the relative impulsivity for food decreased with the willingness to pay for food (*p* = .005; **Fig. 2e**). While slightly counterintuitive, as one could expect that a higher subjective valuation of food should yield more impulsive behaviour, this falls in line with previous results demonstrating that impulse control improves for higher reward prospects ^45^.

Together, these results demonstrate that changes in physiological state induced by fasting modulate impulse control for food rewards. Furthermore, fat reserves moderated these dynamics, suggesting a more complex interplay between short- and long-term energy status in cognitive control. Critically, those effects could not be explained by an increase in the subjective value of food, which, on the contrary, improved performance.

### Relative energy deficit drives motivation to effort

To assess the role of metabolic state in effort regulation, we adapted a classic incentive-motivation task ^46^ (**Fig. 1b**) as our second behavioural measure. Briefly, participants held a dynamometer in their hand, which they could squeeze to raise the level of a thermometer-like scale on the screen, thereby increasing their chances of earning a reward. The colour of the scale indicated the rate at which the thermometer would rise, allowing participants to gauge their behaviour as a function of the effort required to fill the scale (difficulty level) and, therefore, the actual cost/benefit ratio at stake. As for the stop signal task, trials were organised in alternating blocks (indicated by food or money pictures displayed next to the scale), specifying which type of reward would be delivered at the end of the session.

Performances were renormalised to each participant’s strength and analysed using linear models fitted for each cue type and including an intercept, the difficulty level, and trial number. The resulting subject-level estimates were then carried to a group level analysis showing that, unsurprisingly, participants exerted less force and were therefore ready to forego their chances of reward, as difficulty increased (*p* < .001; **Fig. 3a**). While the type of reward at stake also affected the performances (interaction with intercept *p* < .001; difficulty: *p* < .001; trial: *p* = .011), this was mainly driven by the difference in subjective valuation of the outcomes (all *p* < .033) and not by an interaction of metabolic factors (all *p* > .446). Accordingly, we averaged the two conditions before exploring the influence of the energy status on behaviour. Interestingly, fluctuations in average performances were explained by an interaction between fasting duration and body composition (*p* = 0.006, correcting for sex differences, **Fig. 3b**). Indeed, while overall performances drastically declined with body fat percentage (*r* = -.34, *p* = .001, **Fig. 3d**), this effect was partially mitigated by fasting (short fast: 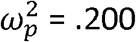; long fast: 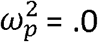; **Fig. 3c**), suggesting that fasting could partly normalise the negative influence of fat mass on motivation. Examining the effect of fasting across subgroups stratified by body fat% (**Fig. 3d**) may help explain this complex pattern. Indeed, motivation appears to be related to fasting when appraised not by its duration but by the relative energy deficit it induces. To test this hypothesis, we estimated the number of calories burned during the fasting period relative to the number of calories stored in the body as fat (see Methods). This measure of relative energy deficit (RED) was strikingly similar to our motivation measure (**Fig. S1**): participants with a high body fat percentage (*i. e*. a slow metabolic rate and large reserves) had a low RED which slowly increased with fasting; in contrast, participants with a lower body fat percentage (*i. e*. burning calories fast, with low energy stocks) had a higher RED which was less consistently affected by fasting as body composition dominated the variations between individuals. Accordingly, RED strongly predicted effort spending (*p* < .001, **Fig. 3e**) and provided a more parsimonious explanation than the fasting × body fat interaction (*Δ* BIC = 2.1).

**Figure 3:**
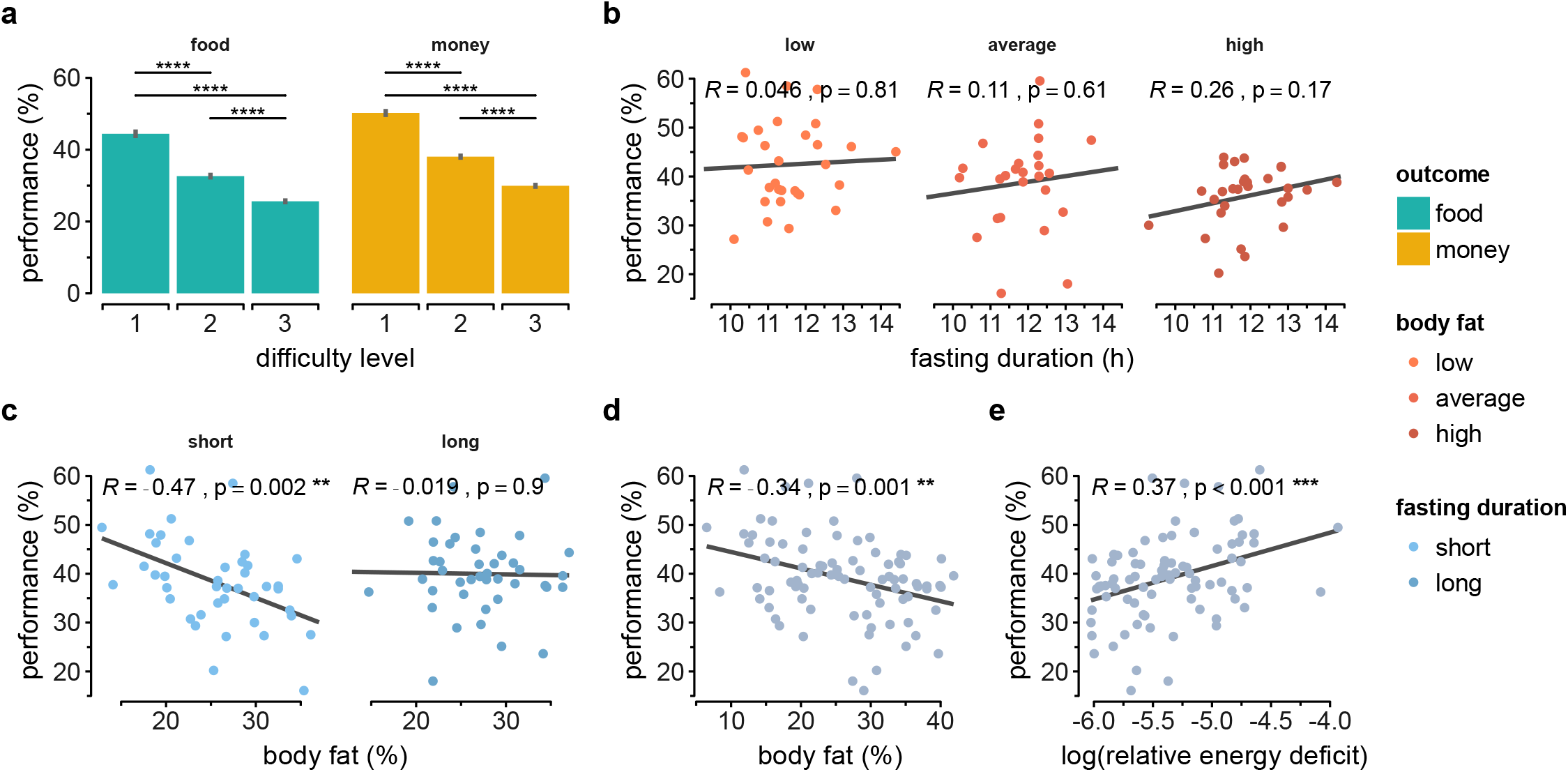
Results of the incentive motivation task. *a)* Average performance (gauge level) for each difficulty level for food (green) and monetary (yellow) rewards. Bar and error bars represent the group average and standard errors; *b)* Effort increases with fasting duration, especially as body fat increases (here split into body fat terciles from left to right: light orange: *≤* 21%, dark orange: >21% and *≤*31%, red: >31%). However, higher body fat was also associated with lower average effort; *c)* This effect of body composition is driven by participants with shorter fasting duration (< 11.5h, left, light blue) and normalises with longer fasting (> 11.5h, right, dark blue); d) Overall, effort decreased with body fat%; *e)* The body composition *X* fasting interaction is summarised as an effect of relative energy deficit induced by fasting. All statistics are Pearson correlations, and the best-fit lines are linear regressions computed for the data shown in each plot.

In summary, our results demonstrate that short-term energy deficits strongly upregulate effort expenditure. In contrast to the impulse control measures reported above, this metabolic effect appears pervasive and does not depend on outcome quantity or identity, both of which also influence motivated behaviour.

### Changes in willingness to pay for food do not explain impulsivity or effort variations

During the first two tasks, capturing impulsivity and motivation, respectively, participants were instructed that good performance would earn them food and money tokens, depending on the block, to be exchanged for actual food items and money at the end of the experiment (**Fig. 1c**). The subsequent auction task was framed as an opportunity to reallocate the tokens they won by betting on 30 pairs of fortune wheels. On a given trial, each of 10 tokens could be placed either on a wheel associated with one of the 30 available food items or on another associated with an equivalent monetary value. As each token granted a 10% chance of the fortune wheel to stop on a win, participants could decide to either place all their bets on one wheel and thus ensure to win the associated outcome or split their bets and have a chance to win both rewards.

The total amount of tokens allocated to snacks, irrespective of the strategy, reflected the willingness to pay (WTP) for food. Although a linear model including fasting duration and body composition did not yield a significant effect, WTP correlated with body fat percentage when tested separately (*p* = .040; **Fig. 4**).

**Figure 4:**
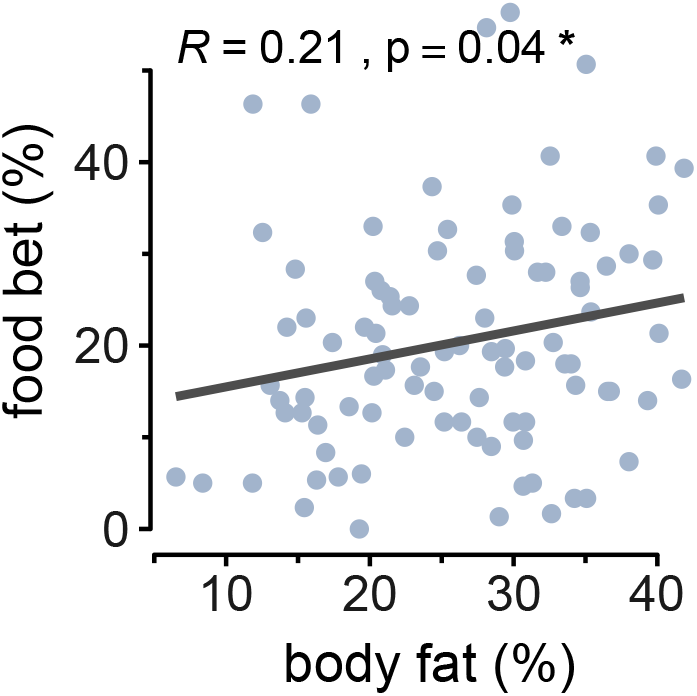
Results of the auction task. Willingness to pay for food increases with body fat percentage.

Importantly, the prospect of reward is a strong predictor of both motivation and impulsivity, with higher incentives associated with better performance on both measures. While WTP for food was not robustly modulated by metabolic state in our data, it could still partially explain the influence of bodily state on performance. To control for this potential confound, WTP was systematically included as a covariate in analyses of the stop-signal and incentive-motivation tasks. However, adding this control did not affect the results.

In conclusion, the influence of metabolic state on behaviour identified above cannot be explained simply by changes in the subjective value of food rewards; rather, it reflects a fundamental regulatory process of action mediated by physiological state.

### Questionnaires capture static, not adaptive, behavioural phenotypes

We performed an exploratory factor analysis to summarise the 15 questionnaires completed by each participant (**Table 2**), yielding five factors (**Fig. 5a**). The first two are related to food behaviour, and more precisely to sensitivity to external food triggers (“uncontrolled eating”, similar to the previously reported factor ^47^, and the active tendency to restrain one’s food behaviour (“cognitive restraint”). The next two factors concern more general behaviour: sensitivity to rewards versus punishment (“drive”) and impulsivity (“impulsiveness”). The last factor captured associations between depressive traits and compulsive tendencies along with food coping strategies (“compulsiveness”).

**Figure 5:**
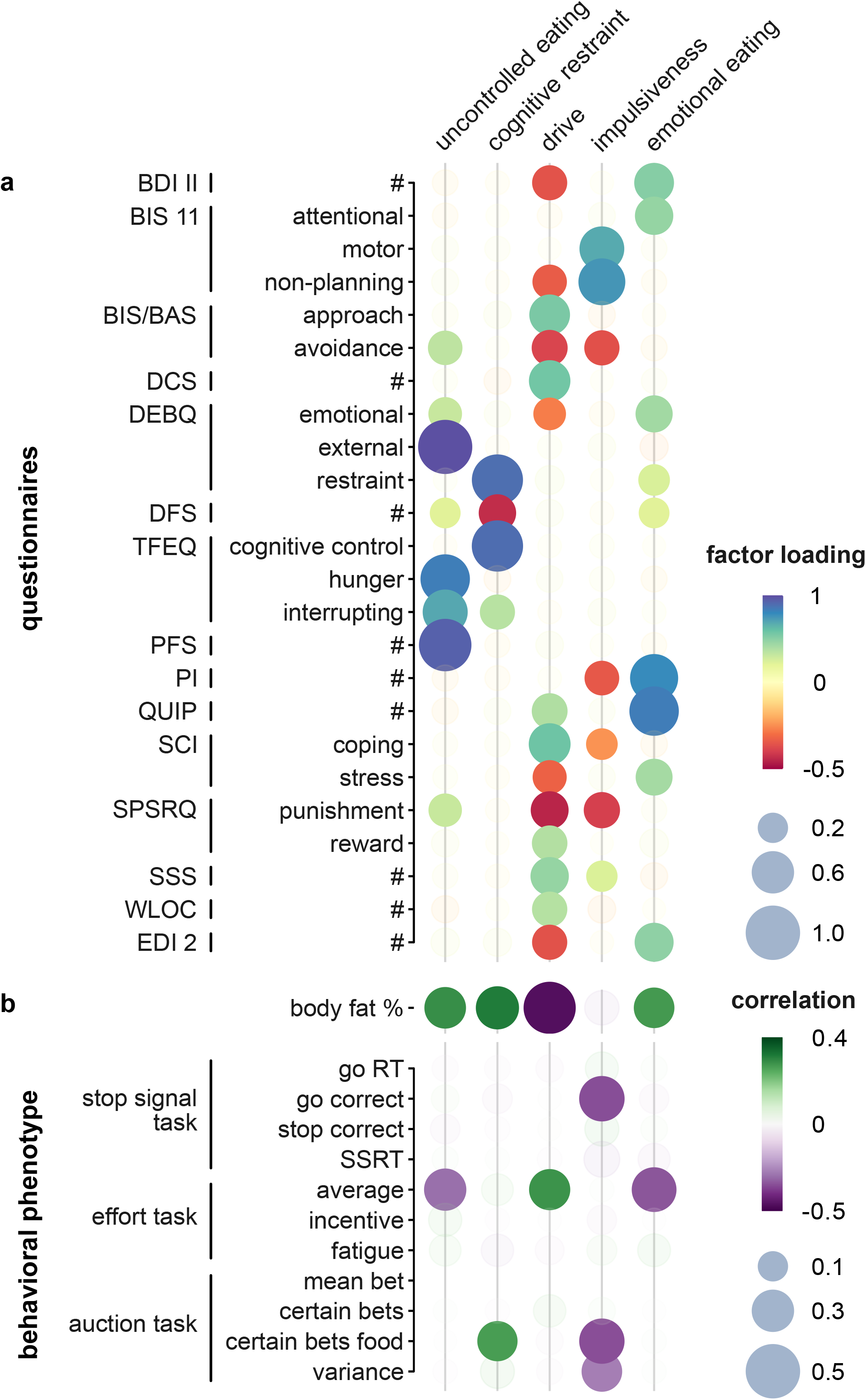
Factor analysis of the questionnaires. *a)* Each dot represents the loading of each questionnaire (sub)scale (rows) on the five identified factors (columns). Factor loadings represent the strength of the association between a scale and a latent factor: larger and darker dots indicate stronger loadings, and colour encodes the sign of the loading (blue: positive, red: negative). *b)* Each dot represents the correlation between factor scores, which capture where each participant lies on the factor space, and empirical measures of body composition or behavioural metrics. Bigger and darker dots represent stronger correlations, and colour encodes loading sign (green: positive, purple: negative).

To understand the link between each participant’s traits, as captured by the self-report questionnaires, and their actual behaviour, we first computed the factor scores, which are a summary measure of where each participant stands on the identified latent factors, and then correlated these factor scores with individual task performances (**Fig. 5b**).

Regarding the stop-signal task, we found a single correlation between accuracy in the go condition (*i. e*., correctly following the arrow direction) and the impulsiveness factor. Although surprising at first, as SSRT is the metric expected to capture impulsivity in this task, this finding is consistent with previous reports showing that inaccurate action responses were more predictive of trait impulsivity than action inhibition ^48^. A more direct explanation for the lack of correlation between trait impulsivity and SSRT performance is that the latter is a highly dynamic phenotype that continuously adapts to physiological state and, therefore, is unlikely to be captured by questions designed to assess static qualities. Overall, our results highlight the fundamental shortcomings of questionnaires when quantifying fast-fluctuating behaviours such as (fasting-dependent, food-specific) impulsivity.

The effort task exhibited a simpler, domain-general behavioural marker, summarised by average effort performance. This metric was positively correlated with the “drive” factor, consistent with the well-known importance of reward sensitivity in regulating effortful actions. Effort was further anti-correlated with the “uncontrolled eating” factor, which could be explained by the fact that this factor also captured a negative drive dimension (*i. e*., sensitivity to punishment), which partially mirrored the “drive” factor. Finally, average effort was negatively associated with the last factor we labelled “compulsiveness”. This factor also loaded on depression and stress questionnaires and could reflect a more general mood downregulation that would negatively affect motivation.

Finally, in the auction task, the proportion of certain bets (the number of snacks secured by placing all tokens on the food wheel) positively correlated with cognitive restraint and negatively with impulsivity factors, suggesting that those traits can translate into risk aversion when implementing actual food choices and are therefore also pertinent to understanding weight regulation.

Next, we explored the link between trait dimensions and body weight. Almost all factors correlated with body composition, indicating that a higher fat percentage is associated with greater sensitivity to food cues, stronger attempts to restrain such urges, and greater compulsiveness. The factors we identified are consistent with previous reports, particularly uncontrolled eating ^47^. Body fat was also associated with lower drive, suggesting that the motivational deficit we measured in the effort task may not be attributable solely to a dampened fasting effect in the more corpulent participants, but also to a more general lack of reward sensitivity.

Interestingly, “impulsiveness” did not correlate with body composition. While various measures of impulsivity have been associated with body weight before, these associations originate from clinical populations suffering from food-related disorders such as morbid obesity or binge eating disorder and might not hold for the general population ^49,50^.

Altogether, our observations show that while self-report questionnaires can capture some behavioural phenotypes and their importance for weight regulation, they overlook the dynamic aspects of behaviour and therefore fail to fully capture the adaptive nature of metabolically regulated behaviour.

## Discussion

In this study, we investigated how short- and long-term energetic states jointly shape adaptive behavioural control in healthy humans. We found that fasting selectively increased food-related impulsivity, whereas motivation to exert effort scaled with relative energetic deficit across reward domains. Importantly, both effects depended on longer-term energy reserves, indicating that identical fasting periods need not represent equivalent physiological challenges. Together, these findings demonstrate that inhibitory control and motivation are governed by distinct metabolic principles operating across multiple energetic timescales. More broadly, these findings provide empirical support for recent theoretical frameworks proposing that behaviour emerges from the predictive, allostatic regulation of physiological state rather than from fixed behavioural traits ^51^.

Regarding impulsivity, our results generalise recent findings identifying a difference in inhibitory control between fed and fasted states in a food-related task ^19^, showing that impulsivity progressively increases with fasting duration. We also demonstrated that the effect of fasting was specific to food rewards, thereby confirming the existence of a domain-specific form of impulsive behaviour ^52^. Notably, another study, using a different measure of impulsivity (information sampling), reported non-food-related changes in impulsivity with hunger ^20^. This discrepancy can be resolved by acknowledging that impulsivity is a multifaceted construct encompassing distinct mechanisms ^53^. In light of this literature, our results suggest that state-dependent modulation of impulsive behaviour may be domain-specific, depending on the underlying process measured. Interestingly, the stop-signal task we used in this study is recognised as yielding highly variable results across individuals ^54^ and showing only modest associations with body composition ^49^. Our findings suggest that at least part of this variability reflects differences in metabolic state, a source of behavioural variation that has rarely been considered in previous studies. Collectively, these results highlight that to understand the exact relation between body composition and impulsivity, future studies should carefully account for the fluctuations (or the lack thereof) induced by the hunger state.

Concerning motivation, our data confirm previous reports showing that the effort to obtain food increases with increasing hunger ^25,55,56^. Notably, those studies relied solely on food rewards, suggesting a food-specific effect. In contrast, our study revealed that fasting increases vigour across outcome types, suggesting a general motivational effect. Importantly, this behavioural effect was better explained by relative energy deficit than by fasting duration itself, indicating that motivation scales with the energetic significance of fasting rather than with time since the last meal per se. This observation suggests that the behavioural impact of fasting depends not on fasting duration alone, but on its physiological significance for the individual. Furthermore, this finding aligns with prior behavioural evidence showing that hunger promotes acquisition-related behaviour beyond food rewards, without altering evaluative preferences, and extends these effects to effort expenditure under explicit energetic constraints ^18^.

We also found a negative correlation between body fat and motivation. While this result also aligns with previous studies ^57,58^, we suggest a different interpretation of this relation: rather than a hallmark of adiposity attributable to dopaminergic dysregulation, the reduction in vigour among participants with higher body fat could be due to a reduced influence of fasting. More precisely, we propose that the behavioural consequences of fasting should be interpreted relative to the organism’s energetic context, defined jointly by immediate energy deficit and longer-term energy reserves. For individuals with greater energy reserves, the same fasting period represents a smaller energetic challenge and therefore exerts a weaker influence on behavioural control. Mechanistically, this could be explained by the observation that, in rodents, the fasting-induced increase in motivation depends on a switch to a ketogenic metabolic regime, the timing of which is determined by fat reserves ^59^. Contrary to this conclusion, other studies have found an increase in motivation with BMI ^24,60,61^. Differences in the experimental design might explain these discrepancies. First, those studies compared lean and obese populations, whereas we tested only healthy-weight participants. Morbid obesity is associated with numerous metabolic and neural changes that could disrupt the normal regulation of motivation by bodily state. Second, they offered high-calorie snack foods, while our selection contained healthy options. As motivation appears to depend on the type of food at stake ^58^, a more granular approach may reveal distinct effects depending on the macronutrient composition of the food reward. Our work underscores the need to account for variations in energy needs to correctly identify motivational phenotypes. More generally, our findings indicate that behavioural responses depend less on absolute physiological state than on the energetic significance of that state for the individual.

The dissociation between inhibitory control and motivation observed here suggests that metabolic signals engage at least partially distinct neural mechanisms governing adaptive behavioural control. Accordingly, our findings are consistent with the metabolic regulation of dopaminergic circuits ^62^, which are critical for the control of willingness to exert effort ^63^ and impulse control ^64,65^. Indeed, hunger-related hormones, such as ghrelin, potentiate dopaminergic activity and thus motivation, while leptin, produced by adipose tissue, tends to dampen it ^30^. Because hormones such as ghrelin and leptin operate on different physiological timescales and convey complementary information about current metabolic state and available energy reserves, they provide plausible substrates through which energetic context could regulate behaviour.

Furthermore, obesity has been robustly associated with dopaminergic dysregulation, although the exact relationship with body weight regulation remains unclear ^66^. A better understanding of how the various metabolic signals interact, at their respective timescales, to control dopaminergic function is needed to decipher state-dependent regulation of behaviour.

From an ecological perspective, our data are consistent with the hypothesis that increased energy requirements shift behaviour toward food acquisition by increasing effort expenditure and biasing decisions toward more immediate actions. While we also found that adiposity was associated with lower motivation, we proposed that this reflects only a blunted or delayed fasting effect in participants with higher energy reserves. Our interpretation is thus at odds with the classical view in the literature, which associates obesity with a set of “traits” or cognitive phenotypes assessed using psychometric questionnaires ^67,68^. The importance of dynamic metabolic regulation, overlooked by this approach, may explain why the association between stable aspects of body composition and cognitive traits is weak ^69^. Our findings suggest that body composition should be viewed not only as the outcome of stable behavioural traits but also as a determinant of how behaviour adapts to changing energetic demands.

Importantly, our conclusions are based on healthy participants exposed to a moderate fasting challenge. Whether similar behavioural dynamics operate under conditions of sustained energy imbalance, prolonged dietary restriction, or clinical obesity remains an open question, as these conditions are accompanied by stable neuroendocrine, inflammatory and neural adaptations that may fundamentally alter behavioural regulation. Future work should examine how metabolic signalling interacts with established neuroendocrine and neural adaptations to shape adaptive behavioural control across different physiological and environmental contexts. Taken together, our findings support a view of adaptive behavioural control as a dynamic process that continuously interprets metabolic signals in the context of available energy reserves to adapt behaviour to physiological need. Understanding how these interacting signals are implemented at the neural level will be an important challenge for future work.

## Methods

### Participants

A total of 116 volunteers were recruited from a local database. A first screening for exclusion criteria allowed us to identify volunteers either being underweight (BMI < 18.5, *n* = 2), obese (BMI > 30, *n* = 6), following a restrictive diet (*n* = 1), having a physiological condition that could affect their food behaviour (*n* = 7), suffering or having a history of psychiatric disorder(s) (*n* = 3), scoring high on depression scale (BDI > 17, *n* = 3), or being pregnant (*n* = 1). One participant was further excluded for not arriving fasted on the day of the experiment. In total, 94 healthy participants underwent the full experimental protocol and were included in the data analysis (**Table 1**).

### Experimental design

Before being invited, participants completed all questionnaires related to food or used for exclusion (**Table 2**) via a dedicated online platform hosted at our institute. They were then asked to refrain from consuming any food or caloric beverages after 10 PM the day before their single testing session, which started at 8 AM. Upon arrival, participants were familiarised with the general course of the experiment. In particular, they were told they would need to perform two behavioural tasks (a stop-signal task and an incentive-motivation task, order counterbalanced across participants), allowing them to earn, depending on the experimental bloc, either “food tokens” or “money tokens”. Participants were informed that the tokens could be exchanged for food items and cash later. Unbeknownst to the participant, the conversion of the collected tokens into actual rewards was carried out using a final “fortune wheel” auction task. This procedure allowed us to measure participants’ willingness to pay for food, *i. e*., to quantify the subjective value of the two reward types relative to each other. Participants were then permitted to consume the food they had won (no other food was allowed) before proceeding with the rest of the experiment, namely completing the paper questionnaires (**Table 2**) and undergoing anthropometric measurements to assess body composition and estimate forearm muscle size.

To ensure that the framing of the tasks in “food” and “money” blocks was clearly mapped onto concrete, distinct outcomes, all available snacks and a cashbox were openly displayed in the room throughout the experiment. In addition, participants rated their liking of each of the 30 food items before starting the behavioural tasks. Participants also rated their levels of hunger, thirst, and satiety throughout the session (seven timepoints).

All procedures were approved by the ethics committee of the University of Cologne (No. 10-226), and we obtained written informed consent from all participants prior to the experiment.

#### Stop signal task

The computerised stop signal task was adapted from the open-source “STOP-IT” software ^44^. Participants rested their index fingers on the left/right arrow keys and looked at a fixation dot on the screen. Following a jittered delay (500-1350 ms), a white arrow appeared (go-signal), and the participant had to indicate the direction of the arrow by pressing the corresponding key (go-trials). On 25% of trials, the arrow turned blue (stop-signal), instructing participants to withhold their response (stop-trials). The trial ended upon response or after 1500 ms (the maximum reaction time).

To assess the influence of the incentive domain, trials were organised in 6 alternating blocks (96 trials each, with a 15-second pause between trials), starting with a full-screen image of the reward at stake (food or money; order counterbalanced across subjects). The same picture remained displayed on both sides of the screen during the trials, flanking the go/stop arrow signal (**Figure 1**). Participants were instructed to be fast and accurate and were informed that their performance in the respective blocks would determine the amount of food and money they would win. A short practice block (32 trials) with textual feedback after each response, but no incentive was included at the beginning of the task.

The stop-signal delay (SSD) was independently adjusted for each reward type based on the participant’s performance using the staircase-tracking procedure i.e., ^70, p. 6892^. The SSD was initially set to 250 ms and adjusted in 50 ms steps based on the response in each stop trial, ensuring that the probability of successful stopping converged to 50 %.

#### Effort task

The effort incentive motivation task is adapted from Pessiglione *et al*. ^46^. Participants held an isometric dynamometer (Vernier Software & Technology) in their dominant hand and were first instructed to squeeze this handgrip as hard as possible (three attempts, 4 s each). The maximal voluntary force (MVF) was then used to calibrate the task’s difficulty.

Each trial started with the display of a thermometer-like scale on the screen. Squeezing the handle (within 3s) filled the thermometer in proportion to the exerted force, reaching the top of the scale, requiring 90 % (easy), 115 % (medium), or 140 % (hard) of the MVF. The difficulty level was indicated by the colour of the “mercury” (respectively green, orange, or dark red), allowing participants to plan their effort before pressing.

As for the stop signal task, trials were organised into 20 domain-specific blocks (12 trials each; order counterbalanced across subjects), starting with a full-screen display of the reward picture, which was also shown on the side of the thermometer for the remainder of the block. Critically, participants were instructed that the higher the mercury, the more tokens, and therefore the more money or food, depending on the cue, they would obtain at the end of the experiment.

#### Auction task

The auction task was run last and allowed participants to trade the tokens they earned for food, snacks, and cash, thereby enabling us to assess both the relative value of different types of outcomes and participants’ attitudes toward risk.

Participants were first informed that they had won 300 tokens during the force and stop-signal tasks and that they now had the opportunity to bid on food and monetary items in a series of lotteries. On each trial, the participant allocated 10 tokens between two independent fortune wheels, one associated with a fixed amount of money (0.70 €), the other with one of the 30 snacks of equivalent value. Each token purchased one segment (1/10th) of the wheel, therefore increasing the probability of winning the corresponding reward. The order and position of the snacks were randomised across participants. After bidding on all lotteries, 6 were implemented by spinning the wheels on the screen and displaying a visual confirmation of the outcome for each wheel. The seemingly random outcomes were biased to ensure that the participant won their three most desired snacks, thereby allowing observation of subsequent food consumption during the questionnaires.

#### Hedonic ratings

Participants were presented pictures of each snack with the question “How strongly do you like or dislike this item?” (“Wie stark ist Ihre Vorliebe bzw. Abneigung”) and rated their preference on a labelled hedonic scale (Lim et al., 2009) ranging from “greatest imaginable dislike” (“Stärkste Abneigung, die vorstellbar ist”) at the bottom to “greatest imaginable like” (“Stärkste Vorliebe, die vorstellbar ist”) at the top.

#### State ratings

Subjective state (hunger, satiety, and thirst) was measured using a visual analogue scale. For each dimension, a question on the screen (“How hungry/sated/thirsty are you at the moment?”; “Wie hungrig/satt/durstig sind Sie momentan?”) prompted the participant to rate their current state between “not hungry/sated/thirsty at all” (“gar nicht hungrig/satt/durstig”) and “very hungry/sated/thirsty” (“sehr hungrig/satt/durstig”) anchors.

#### Anthropometric measurements

Body composition was measured using a SECA mBCA 515/514 bioelectrical impedance analysis scale, which, in addition to total body weight, provided absolute fat mass, fat percentage, fat-free mass, and skeletal muscle mass (full body and limb by limb). We also measured participants’ height to calculate the body mass index (BMI). Circumference, length, and skinfold fat of the forearm were also measured to derive an approximation of the muscle size.

#### Software

All tasks were run using Psychtoolbox 3.0 (http://psychtoolbox.org) on MATLAB (The MathWorks, Inc.). The measurements from the hand dynamometer were captured using a homemade Matlab code (https://github.com/lionel-rigoux/verniertoolbox).

### Statistical analyses

Behavioural and physiological data were preprocessed in MATLAB (v2023a; The Mathworks Inc.) and loaded in R (v4.3.0; R Foundation for Statistical Computing) for analysis using linear (package ‘stats’ v4.3.0) and mixed-effects (package ‘lmerTest’ v.3.1.3; doi:10.32614/CRAN.package.lmerTest) models. Significance was then computed using type III ANOVAs (‘car’ v3.1.2 and ‘lmerTest’ packages, respectively). Correlations were computed using Pearson’s correlation coefficient. For all statistical tests, the significance threshold was set at α = .05. To assess the general or relative domain specificity of the effects, behavioural metrics were either averaged across incentive conditions or subtracted, using *Δ* “food” minus “money”. To illustrate interaction effects, fasting duration and fat percentage were categorised into quantiles, allowing visualisation and estimation of effects within each subgroup; however, statistical tests of interaction were systematically performed on the full, continuous variables.

#### Stop signal reaction time

For each participant and incentive condition, reaction times in go trials were summarised using the geometric mean to reduce the influence of outliers. Stop-signal reaction time (SSRT) was estimated using a logistic regression of stopping accuracy as a function of stop-signal delay (SSD). The SSD corresponding to a 50% probability of successful stopping was identified and subtracted from the mean go reaction time to yield the SSRT (see Supplementary Methods). We excluded from further analyses participants whose performances were too low (go-trial error rates exceeded 10% or omission rates exceeded 25%) or whose stopping accuracy fell outside the 40–60% range, indicating failure of staircase convergence.

#### Effort task

For each trial, peak grip force was extracted from the force profile and rescaled to the maximal physiological force computed from forearm measurements (see Supplementary Method) to obtain a normalised performance (in % of MPF). Within each participant and incentive condition, performance was then modelled as a function of trial difficulty and trial number using multivariate linear regression, yielding estimates of average effort expenditure and sensitivity to task demands. Subject-level estimates were then entered into group-level linear models to assess the influence of metabolic variables and body composition.

#### Auction task

For each lottery, we computed the average proportion of tokens allocated to food outcomes as the willingness to pay for food (WTP. While this means betting indexed preference for food relative to money, the variance in the allocation pattern indexed the betting strategy. To obtain a preference-independent measure of risk aversion, observed variance was normalised by the maximal theoretical variance given each participant’s mean preference (see Supplementary Methods).

#### Physiological measures

Relative energy deficit (RED) was computed as the ratio of estimated caloric expenditure during the fasting period, based on basal metabolic rate, to total energy stored in body fat. This measure captures the energetic cost of fasting relative to long-term energy reserves.

The maximal physiological force (MPF) was computed to be proportional to the physiological cross-sectional area of the forearm muscles, which we derived from anthropometric measurements.

See Supplementary Methods for the complete derivation of those measures.

## Data Availability

All data are available upon request. The request requires that the purpose of data reanalysis align with the study aims, as approved by the ethics review board (see text), and that participants’ consent be obtained. Furthermore, consent to the processing of personal data must be obtained through a signed Material and Data Agreement.

## Acknowledgements

The authors are extremely grateful to Katharina Rüth (neé Schmitz) for tenaciously pursuing data acquisition. *Funding:* The study was supported by the Deutsche Forschungsgemeinschaft (DFG; German Research Foundation) under Germany’s Excellence Strategy – EXEC 2030–390661388 and through SFB 1451, project ID 431549029, C06, as well as by the German Centre for Diabetes Research, project ID 82DZD05H1G (MT).

## Author Contributions

All authors contributed to the work presented in this paper. MT, KES, and LR conceptualised the study; LR and BK designed the experimental setup; LR also organised data acquisition and performed statistical analyses, with support from CM; the manuscript was written by LR and MT and edited by all remaining authors. The required infrastructure was provided, configured, and managed by MT and FJ. The study was supervised by MT.

## Competing Interests

The authors declare no competing interests.

## Notes

### Competing Interest Statement

The authors have declared no competing interest.

### Summary of Updates

This version includes editorial revisions to improve the framing and presentation of the work. The title, abstract, introduction, and discussion have been revised to better emphasize the biological context and significance of the findings. No new data or analyses have been added, and the results and conclusions remain unchanged.

