## Supplementary Materials for "Metabolic state and energy reserves jointly regulate adaptive behavioural control"

### Supplementary Material

**Supplementary Figure**

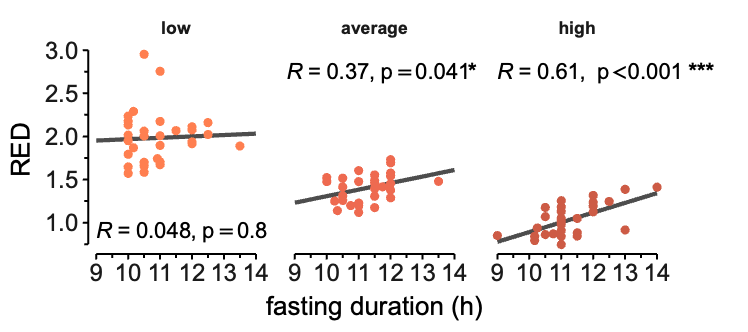

Figure S1: **Relative energy deficit.** Relative energy deficit as a function of fasting duration for the low (left), average (middle), and high (right) body fat participants.

**Supplementary Methods**

***Computation of the stop signal reaction time (SSRT) in the stop-signal task***

We first computed the average reaction time (RT) in the GO condition ($RT_{\mathrm{GO}}$) as an indicator of response process latency. We used a geometric mean to counterbalance the heavy tail of the RT distribution and thereby avoid overestimation. To obtain a signature of the relative efficiency of the inhibition process, we estimated the Stop Signal Reaction Time (SSRT, cf. Matzke et al., 2018 for a review): First, we identified the stop signal delay (SSD) for which the participant had a 50% chance of correctly stopping, known as the critical SSD ($\mathrm{SS}D_{\mathrm{crit}}$), which reflect the average latency of the inhibitory processes. To this end, we performed a logistic regression to predict all the responses in the STOP condition as a function of the SSD, and then located the inflection point. Finally, the SSRT was computed as the difference between inhibitory and response latencies $SSRT=SSD_{\mathrm{crit}}-RT_{\mathrm{GO}}$. Note that a shorter SSRT corresponds to a relatively faster suppression of the action and, therefore, to a more efficient inhibitory process.

**Estimation of risk aversion in the auction task**

In the auction task, participants split their tokens between food and monetary lotteries. While the proportion ($p_{t}$, between 0 and 1) of tokens spent on the food wheel in a given trial reflects, on average, the general preference for the snacks, the variance of the betting pattern across trials ($\mathrm{var}(p_{t})$) is also reflecting, although indirectly, the risk attitude of the participant. Let’s take the example of a participant with a slight preference for food, with an average response $\left\langle p_{t} \right\rangle=0.6$. One strategy to implement this preference is to always split the bet proportionally to the preference: in our example, this would mean always betting 6 tokens on food and 4 tokens on money ($p_{t}$ = 0.6). This strategy is the riskiest, as there is an equally large chance of winning or losing all rewards. Another strategy is to bet, on each trial, all tokens on one wheel ($p_{t}$ = 0 or 1), and to balance which reward type is secured across trials. In our example, this would mean going all-in on the food wheel in 18 trials (60% of 30) and on the money wheel in the remaining 12 lotteries. In that case, the outcome is completely certain.

In the first, risky strategy, the inter-trial variance will be minimal $var\left( p_{t} \right)=0$. In the second, safe strategy, the variance is maximal with $var\left( p_{t} \right)$ = $\left\langle p_{t} \right\rangle\left( 1-\left\langle p_{t} \right\rangle\right)$. By dividing the measured response variance by this maximal theoretical variance, we obtain a standardised (between 0 and 1) measure of risk aversion:

$risk aversion= \frac{var(p_{t})}{\left\langle p_{t} \right\rangle\left( 1-\left\langle p_{t} \right\rangle\right)}$.

**Estimation of maximal physiological force (MPF)**

In the effort motivation task, performance is initially calibrated to the maximal voluntary force (MVF). However, because it is itself a motivated behaviour, the MVF could be affected by fluctuations in energy levels and thus be a potential confounding factor. To circumvent this problem while accounting for interindividual differences in maximal force-generating capacity, we renormalised the effort performances to an objective estimate of the participant's maximal physiological force (MPF), derived solely from anthropometric measurements.

The maximal force a muscle can generate is indeed directly proportional to the number of fibres it contains, which can be approximated by the muscle’s physiological cross-sectional area (PCSA; Maughan et al., 1983) — in our case, the muscles of the forearm in charge of gripping. Practically, we first measured the length ($L$, between the ulna’s head and the styloid process) and maximum circumference ($C$, 1/3rd from the ulna’s head) of the forearm. Then, we used callipers to measure the skinfold ($S,$skin + fat layers) of the interior and exterior sides of the forearm. Approximating the forearm geometry as a cylinder, the PCSA can then be computed as the total area of the limb section minus the fat + bone area (Heymsfield et al., 1982):

$$PCSA=\frac{\left( C-\pi S \right)^{2}}{4\pi}-B$$

where $C$ and $S$ are in cm, and $B$ the bone area, in cm${}^{2}$ (set to 1.8 according to Hsu et al. (1993)). Finally, the MPF can be calculated by simply scaling the CSA:

$$MPF=PCSA\times F$$

where $F=2.45+0.288 \times L$ was previously measured in a large cohort of adults (Neu et al., 2002).

We validated our measure by regressing the MPF on the body composition measures. As expected, the MPF was strongly predicted by participants’ fat-free mass, which is mainly composed of skeletal muscle (main effect: p < 0.001). Critically, this relation was not affected by the fat mass (interaction term: p = .398) nor, alternatively, by the fat mass percentage (interaction term: p = .545).

**Relative Energy Deficit (RED)**

As individuals burn through their energy reserves at different rates, fasting may not have the same effect on all participants. To estimate the relative energy deficit (RED) induced by fasting, we calculated the ratio of estimated energy expenditure during the fasting period to the energy available in the body, stored in adipose tissue.

To obtain the caloric deficit induced by fasting, we first calculated the basal metabolic rate (BMR), which represents the number of calories burned by the body at rest in a day, using the Mifflin-St Jeor equation (Mifflin et al., 1990):

$$BMR = 370 + (21.6 \times LM)$$

where $LM$ is the lean mass measured with the impedance scale. The energy deficit ($E_{deficit})$ is obtained by simply rescaling the BMR to the duration of the fast (T, in hours):

$E_{deficit} = BMR \times T/24$.

The energy stored in the body $E_{available}$ can be obtained directly from the absolute fat mass ($FM$) measured with the impedance scale, assuming that a gram of body fat stores around 9 kcal:

$E_{available} = 9000 \times FM$.

Finally, the RED can be formally expressed as the ratio between the caloric deficit induced by fasting and the calories available in the fat storage:

$RED = E_{deficit} / E_{available}$.

**Exploratory factor analysis**

All questionnaires’ scores were first standardized (z-scored). For questionnaires with multiple subscales, scores for the subscales were used in place of the total score. We then checked that all scales had a Kaiser-Meyer-Olkin factor adequacy larger than 0.5, ensuring enough shared variance amongst questionnaires. Finally, exploratory factor analysis (EFA) was conducted using maximum likelihood estimation. The number of factors was determined by comparing the observed eigenvalues with those of random data in a parallel analysis (function 'fa.parallel' from package 'psych' v2.3.3). To allow for correlation between them, factors were extracted using promax oblique rotation, and score were computed using the ten Berge method (function 'fa.fa' from package 'psych').
